# Evolutionary dynamics of *DIRS-like* and *Ngaro-like* retrotransposons in *Xenopus laevis* and *Xenopus tropicalis* genomes

**DOI:** 10.1101/2021.08.05.455354

**Authors:** Camilla Borges Gazolla, Adriana Ludwig, Joana de Moura Gama, Daniel Pacheco Bruschi

## Abstract

Anuran genomes have a large number and diversity of transposable elements, but are little explored, mainly in relation to their molecular structure and evolutionary dynamics. Here, we investigated the retrotransposons containing tyrosine recombinase (YR) (order DIRS) in the genome of *Xenopus tropicalis* and *Xenopus laevis*. These anurans show 2n = 20 and the 2n = 36 karyotypes, respectively. They diverged about 48 million years ago (mya) and *X. laevis* had an allotetraploid origin (around 17-18 mya). Our investigation is based on the analysis of the molecular structure and the phylogenetic relationships of 95 DIRS families of *Xenopus* belonging to *DIRS-like* and *Ngaro-like* superfamilies. We were able to identify molecular signatures in the 5’ and 3’ non-coding terminal regions, preserved open reading frames (ORFs) and conserved domains that are specific to distinguish each superfamily. We recognize two ancient amplification waves of *DIRS-like* elements that occurred in the ancestor of both species and a higher density of the old/degenerate copies detected in the *X. laevis. X. tropicalis* showed more recent amplification waves estimated around 16 mya and 3.2 mya and corroborate with high diversity-level of families in this species and with transcriptional activity evidence. *Ngaro-like* elements presented less diversity and quantity in the genomes, although potentially active copies were also found. Our findings highlight a differential diversity-level and evolutionary dynamics of the YR retrotransposons in the diploid *X. tropicalis* and *X. laevis* species expanding our comprehension of the behavior of these elements in both genomes during the diversification process.

## INTRODUCTION

Transposable elements (TEs) are the most variable feature of the vertebrate genome, and their role in shaping genomic diversity has attracted considerable interest in recent years (Bourque et al. 2018; Wicker et al. 2018). An exceptional diversity of TEs has been reported in all the amphibian genomes sequenced so far (Hellsten et al. 2010; Sun et al. 2012; Jiang et al. 2015; Session et al. 2016; Hammond et al. 2017; Edwards et al. 2018; Seidl et al. 2019). In *Xenopus tropicalis*, a model organism for genomic studies, TEs represent approximately one-third of the genome (Hellsten et al. 2010). Despite the considerable abundance of TEs in genome annotation, the diversity, molecular structure, and evolutionary dynamics of these elements are still poorly understood. The DIRS elements are a good example of this richness that has not been explored.

Retrotransposons of the order DIRS are widely distributed in the eukaryote genomes (Wicker et al. 2007), except for the birds and mammals (Poulter and Butler 2015). The unifying feature of these elements is that they encode a tyrosine recombinase (YR) which participates in the process of integrating the element into the genome (Poulter and Butler 2015). Other retrotransposons employ endonucleases (LINEs and PLEs) or DDE-type integrase (LTRs) (Wicker et al. 2007).

The DIRS elements were named in recognition of the first retrotransposon containing YR to be described, *DIRS-1*, which was found in the slime mold *Dictyostelium discoideum* (Cappello et al. 1985). This order can be divided into four superfamilies based on sequence structure and phylogeny: *DIRS-like, Ngaro-like, PAT-like*, and *VIPER-like* (Ribeiro et al. 2019). In general, the DIRS elements have three open reading frames (ORFs). The first ORF corresponds to a gag-like domain, the second corresponds to the reverse transcriptase (RT) and RNAse H (RH), and the third corresponds to the YR. Another characteristic of these elements is that the ORFs frequently overlap and have terminal repeats that vary in structure among the superfamilies (Poulter and Goodwin 2005; Ribeiro et al. 2019).

The *DIRS-like* elements can present a conserved methyltransferase (MT) domain downstream from the RT/RH, although the function of this domain is still unknown (Goodwin et al. 2004; Poulter and Butler 2015). The non-coding portion varies in its sequence among the elements, although its basic structure is composed of inverted terminal repeats (ITRs) and an internal complementary region (ICR), which is complementary to the beginning of ITR5’ and the end of the ITR3’ (Cappello et al. 1985; Poulter and Butler 2015).

The *Ngaro-like* elements were described after the *DIRS-like* and are distinguished by their split direct repeats (SDR), composed of A1 in the 5’ end and B1, A2, and B2 in the 3’ end, where A1 and A2 are identical, as are B1 and B2 (Goodwin et al. 2004). These elements do not contain the MT-like domain found in the *DIRS-like*, although in amphibians, they contain an ORF encoding a hydrolase domain (Hydro - SGNH) after the YR, but with no proven function (Goodwin and Poulter 2004; Poulter and Butler 2015).

The *PAT-like* elements are phylogenetically closely related to the *DIRS-like* elements (Goodwin and Poulter 2001; Goodwin and Poulter 2004; Poulter and Goodwin 2005), although these two groups are not always monophyletic (Ribeiro et al. 2019), and they can be differentiated by structural differences in the terminal repeats. Such as for *Ngaro-like, PAT-like* elements are composed of SDRs (Poulter and Goodwin 2005; Ribeiro et al. 2019). The *VIPER-like* elements also have SDRs, and form a distinct group of retrotransposons restricted to the protozoans of the order Kinetoplastida (Ribeiro et al. 2019).

In the Anura, both *DIRS-like* and *Ngaro-like* elements have been described in *X. tropicalis* and *X. laevis* (Goodwin et al. 2004; Hellsten et al. 2010; Poulter and Butler 2015). These species are found across sub-Saharan Africa and have an aquatic life that distinguishes them from other anurans (Hellsten et al. 2010). The *X. tropicalis* karyotype is composed of 2n=20 chromosomes with an estimated genome size of 1.7 Gbp (Hellsten et al. 2010) while the *X. laevis* karyotype has a diploid number of 2n=36 chromosomes, which originated from a process of allopolyploidy, with an estimated size of 3.1 Gbp (Session et al. 2016). The available estimates indicate that 1% of the *X. tropicalis* genome is composed of distinct families of DIRS, some of which may still be active (Hellsten et al. 2010). Evidence of the transcriptional activity of the DIRS elements has already been found in both species (Poulter and Butler 2015), which highlights the possible role of these elements in genome function and evolution.

In the present study, we evaluated the diversity, molecular structure and evolutionary dynamics of the elements of the order DIRS in *X. tropicalis* and *X. laevis*. We identified the structural characteristics of the YR retrotransposons of the DIRS order in both genomes and described diagnostic characteristics for the best differentiation of the elements of the *DIRS-like and Ngaro-like* superfamilies and evaluated the evolutionary dynamics of these superfamilies in these genomes.

## MATERIAL AND METHODS

An extract containing all the elements identified as DIRS was obtained from the Rebpase database (Jurka 2000) version 23.11. All the sequences from *X. tropicalis* and *X. laevis* were selected and analyzed using the NCBI “*Open Reading Frame Finder*” (ORFfinder) (https://www.ncbi.nlm.nih.gov/orffinder/) to identify ORFs with default parameters (“minimal ORF length (nt)” = 75; “Genetic code”: 1. Standard; “ORF start codon to use”: ATG only). The presence of conserved domains was analyzed using the NCBI “*Conserved Domains Search Service*” (CD-Search) (Marchler-Bauer and Bryant 2004) with an e-value threshold adjusted to 0.1. The presence of inverted terminal repeats (ITRs), internal complementary region (ICRs), and split direct repeats (SDRs) was investigated using NCBI BLASTn with the same sequence as query and subject, selecting the options “Align two or more sequences” and “somewhat similar sequences (blastn)”, with the “word size” parameter being adjusted to the minimum and the evalue threshold was 10.

Three families of *X. tropicalis* were selected for the analysis of copies in the genome, including one *DIRS-like* family (DIRS-37_XT) and two *Ngaro-like* families (DIRS-53_XT and DIRS-54_XT). We chose the DIRS-37_XT and DIRS-53_XT families as queries because they have the conserved structure of the ORFs and the complete domains, as well as the characteristic repeats for each superfamily. The DIRS-54_XT family was also used as a query to expand the searches of *Ngaro-like* even despite not having conserved terminal repeats.

The amino acid sequences corresponding to the RTs of both elements were used as queries in online tBLASTn searches against the *X. tropicalis* (GCA_000004195.4) and *X. laevis* (GCA_001663975.1) genomes. The first ten hits were retrieved with 3 kb of both upstream and downstream regions. All the copies retrieved were analyzed for the identification of the ORFs, the conserved domains, and the repetitive regions as described above.

The evolutionary analyses were based on the alignment of the RT amino acid sequences, including the following sequences: (i) the consensus sequences of the DIRS families of *X. laevis* and *X. tropicalis* recovered from Rebpase; (ii) the copies that are homologous to the DIRS-37_XT, DIRS-53_XT, and DIRS-54_XT families retrieved from the *X. tropicalis* and *X. laevis* genomes; and (iii) elements known to belong to the different superfamilies of the order DIRS, *Ngaro-like* - Ngaro1_DR (AY152729 - *Danio rerio*) and Lv_Ngaro2 (AGCV01398517 - *Lytechinus variegatus*), *PAT-like* - SkowPAT (Rebpase - *Saccoglossus kowalevskii*), and *PAT* (Q26106 - *Panagrellus redivivus*), and *DIRS-like* - DIRS-1_Acar (Rebpase - *Anolis carolinensis*) and DIRS-5_CBP (Rebpase - *Chrysemys picta bellii*).

The sequences were aligned using the PSI-coffee tool (Notredame et al. 2000), with Genedoc 2.7 (Nicholas *et al*., 1997) being used for sequence manipulation and editing. Most of the RT sequences contained around 200 amino acids (aa), and sequences with less than 70% coverage were excluded from the matrix. The MegaX program (Kumar et al. 2018) was used to determine the best amino acid substitution model. The phylogenetic tree was reconstructed using Bayesian inference (BI), run in MrBayes 3.2.6 (Ronquist et al. 2012) based on the LG+G model. The Markov Chain Monte Carlo (MCMC) was run for 10,000,000 generations, sampled every 1000 generations, with 25% of the initial results being discarded as burn-in. The final tree was visualized and edited using iTOL (Letunic and Bork 2019).

The copies were named *a priori* according to the family used as the query, abbreviated to D37, D53 and D54, followed by the number of the copy referring to the order in which the sequence was recovered, while “XT” and “XL” are acronyms for *X. tropicalis* and *X. laevis*, respectively.

For the landscape analysis, the consensus sequences available in the Repbase for each *DIRS-like* family of *X. tropicalis* and *X. laevis* and each each *Ngaro-like* family of *X. tropicalis* were used to compose the *DIRS-like* and *Ngaro-like* sequence libraries of each species. In the case of the *Ngaro-like* elements of *X. laevis*, as the families were not available in the Rebpase, the consensus sequences were obtained from the copies recovered in the genomic search described above. For that, the sequences were aligned using MAFFT v7.471, with the pairwise divergence being assessed using Genedoc 2.7 (Nicholas et al., 1997) separating the sequences with more than 80% divergence into distinct groups (considering as different families). The consensus sequence of each group was obtained using UGENE (simple extended algorithm) with a 50% threshold (Okonechnikov et al. 2012). The *DIRS-like* and *Ngaro-like* libraries of each species were used to screen the genomes using RepeatMasker 4.1.0 (with the “-s”, “-nolow “, “-no_is”, “-a”, and “-lib” options). RepeatMasker utility Perl scripts were used to summarize the output (script buildSummary.pl) and to calculate Kimura 2-Parameter (K) divergence with adjusted CpG (script calcDivergenceFromAlign.pl). The scatter plot graphs representing the repeat landscape were created using the Python Matplotlib-v3.3.2 (Hunter, J.D., 2007) and edited in Inkscape software.

The age of the copies was estimated based on the time since the divergence of the ancestral sequence (since the consensus of each family used in the RepeatMasker is an approximation of its ancestor) using the formula: T = K / r (Jiang et al. 2002), where a divergence (K) was obtained as described above, and r is the nucleotide substitution rate of 3.1 x 10^-9^ substitutions per year, which is the average of the estimated substitution rates for *X. tropicalis* and *X. laevis*, and for the L and S subgenera of *X. laevis* (Session et al. 2016).

In order to investigate which families are being expressed in *X. tropicalis* and *X. laevis*, the expressed sequence tags (EST) library from both species were retrieved from Xenbase (http://www.xenbase.org/, RRID:SCR_003280) and the different DIRS families of both species were used as queries in BLASTn. The results were filtered by identity (>85%) and size (>100 bp).

## RESULTS

### Xenopus DIRS sequences belong to DIRS-like and Ngaro-like families

A total of 75 YR-retroelement families were identified in the Rebpase for *X. tropicalis* and 20 for *X. laevis* (Table 1). In this database, all YR-containing elements are classified as DIRS, a final group within the long terminal repeat (LTR) retrotransposons group. We thus previously assume that these families could belong to any of the DIRS superfamilies, then our analyses indicate they belong only to *DIRS-like* and *Ngaro-like*.

**Table 1:**
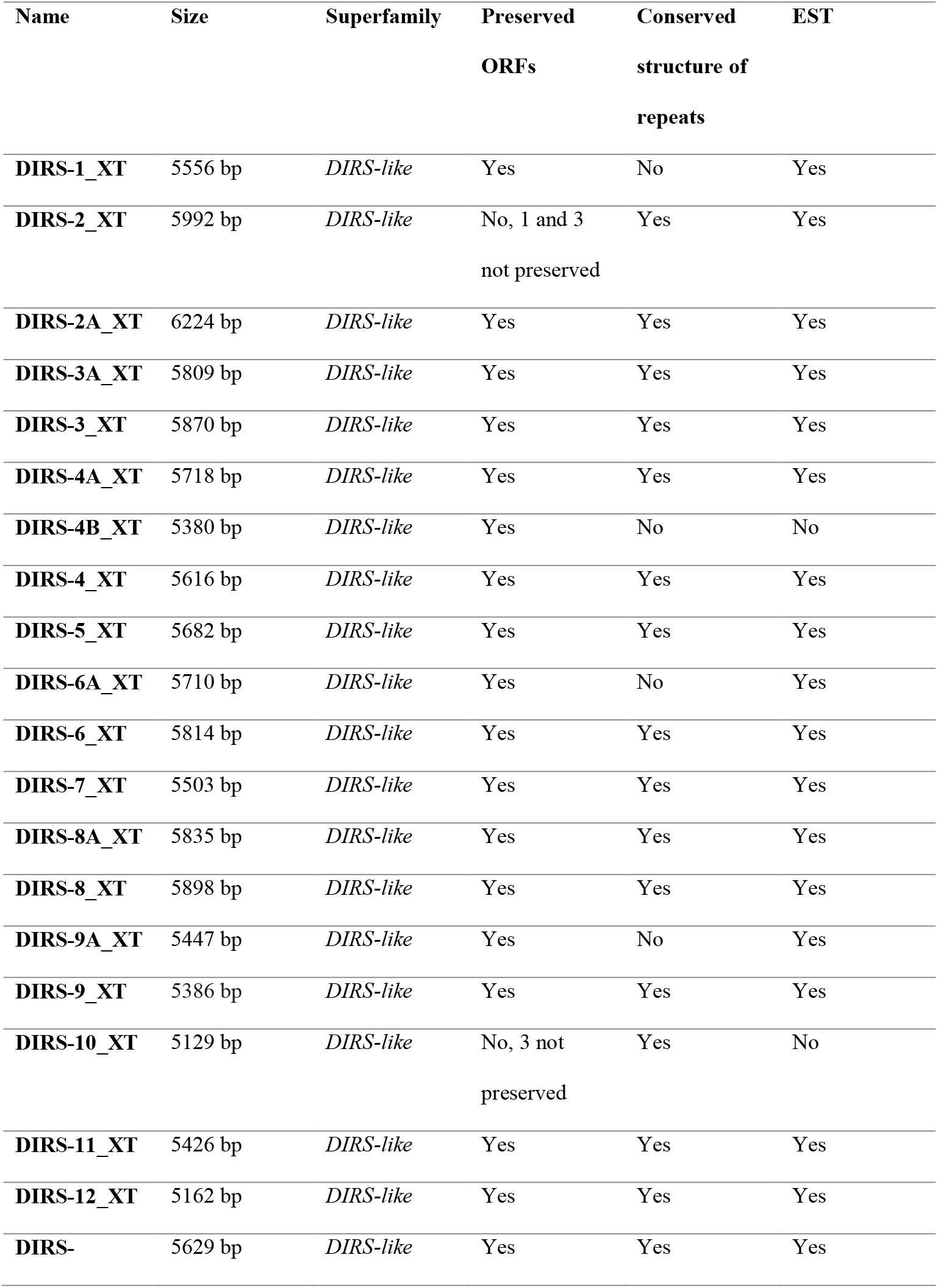
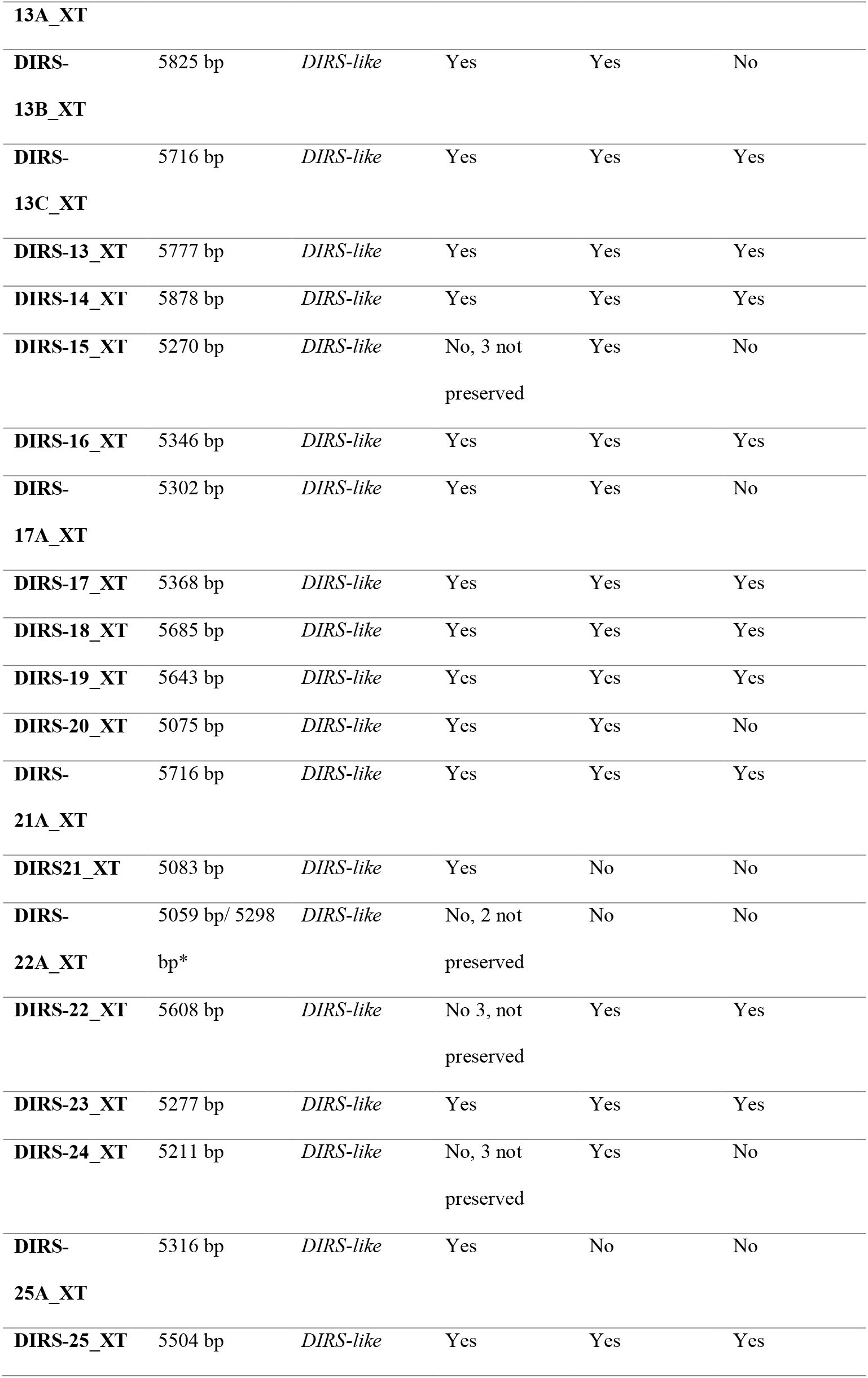
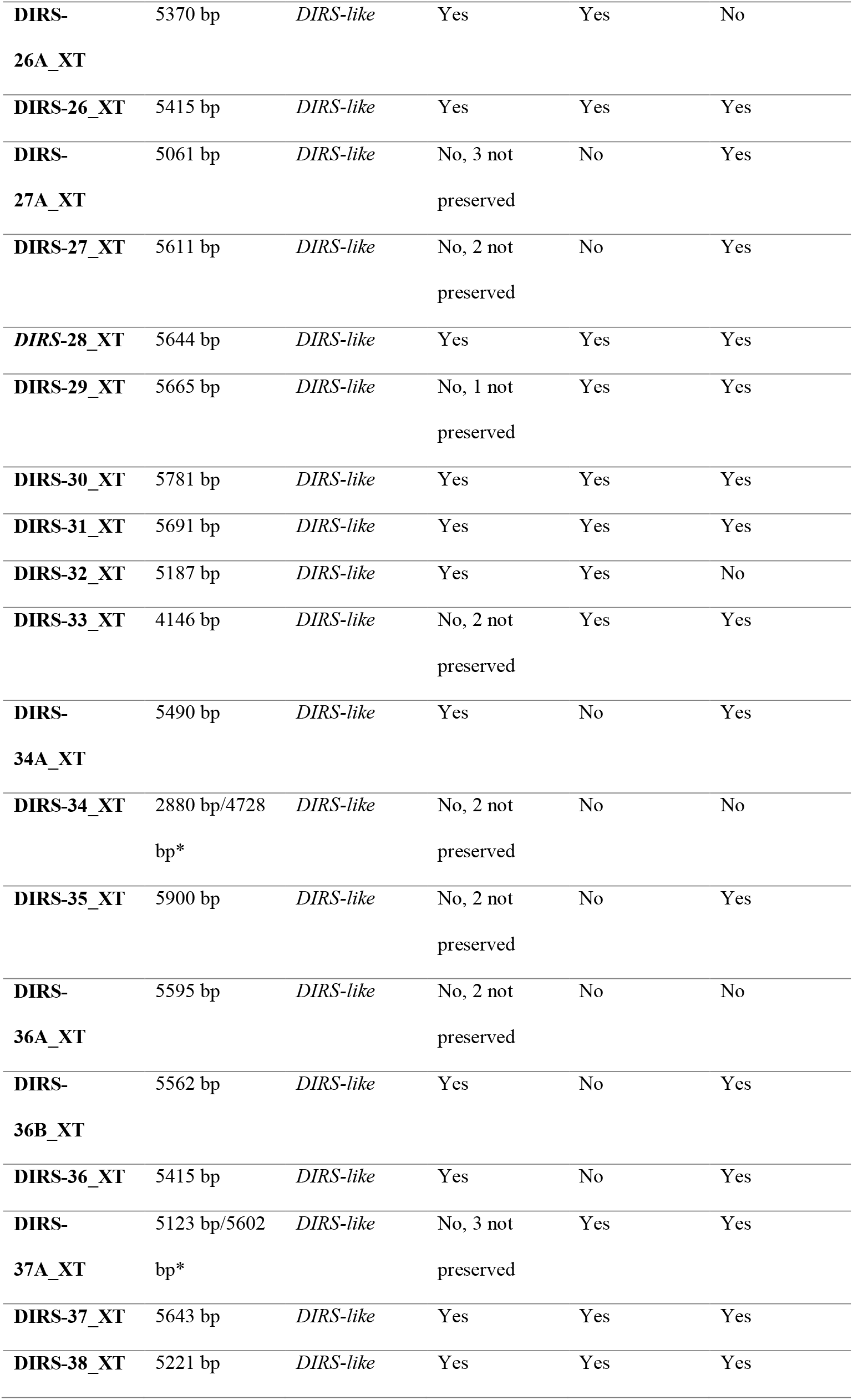
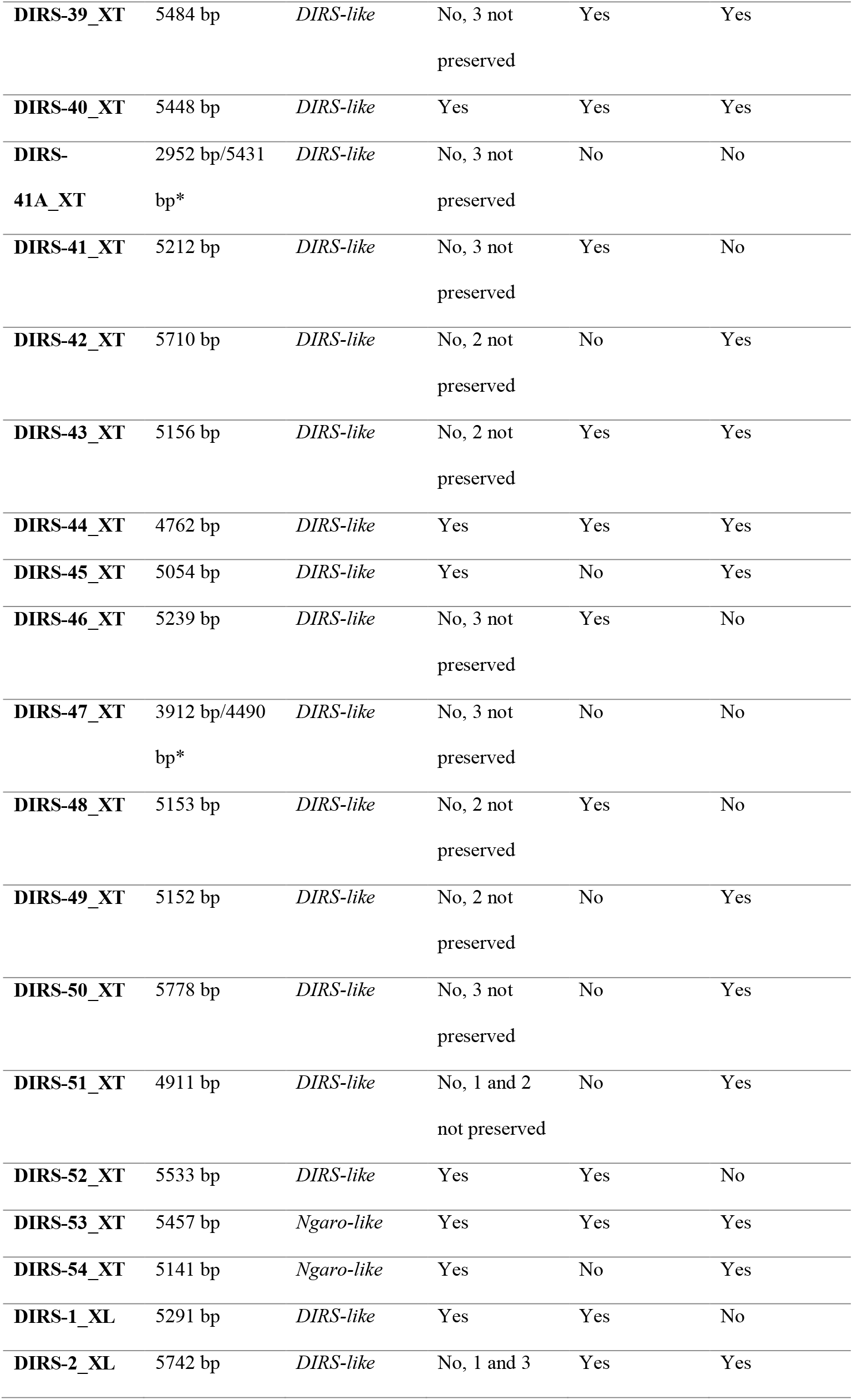
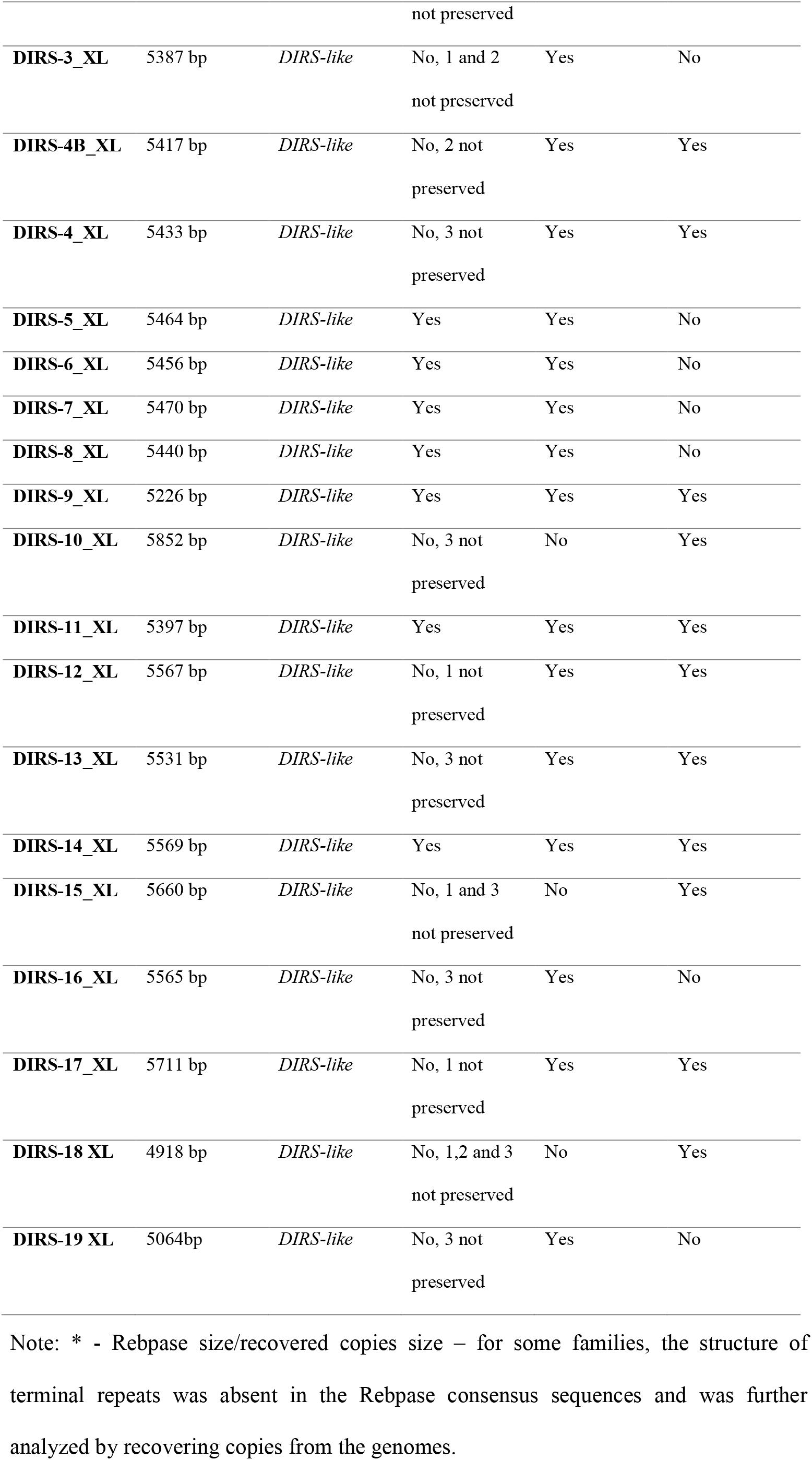
Basic features of the DIRS families of *Xenopus tropicalis* (XT) and *X. laevis* (XL) deposited in Rebpase. The consensus sequence of each family was evaluated for the presence/absence of complete conserved ORFs containing the expected domains and the presence of complete structure of repeats (ITRs and ICR for *DIRS-like* and SDRs for *Ngaro-like*). The presence of ESTs is also shown for each family.

The evolutionary tree based on the RT domain of all the elements together recovered two well-supported clades (Figure 1) in which all the *Xenopus* sequences grouped in either (*i*) a *DIRS-like* or (*ii*) a *Ngaro-like* group. The two *PAT-like* sequences were not grouped as a monophyletic group (Figure 1).

**Fig. 1.**
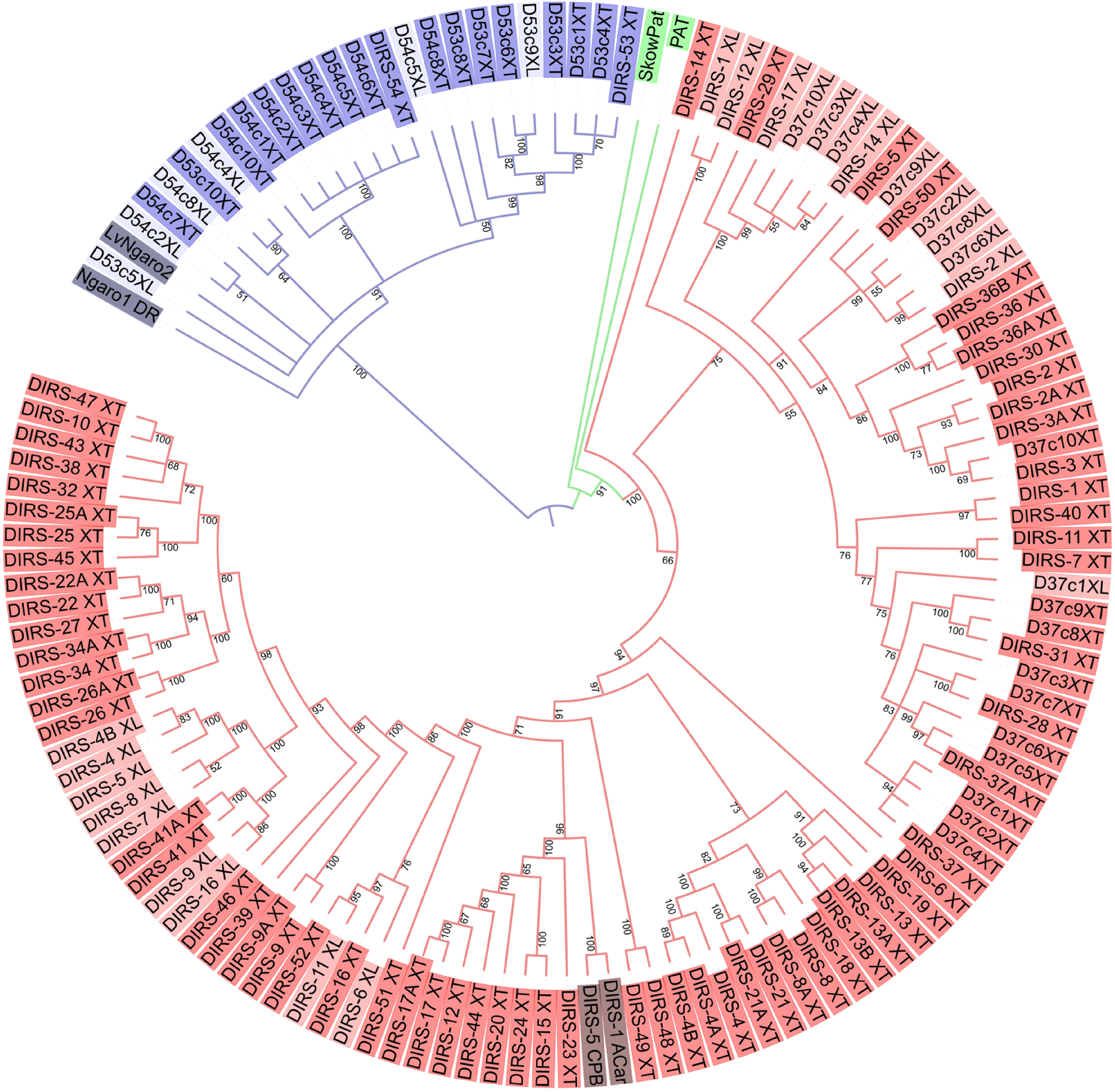
Sequence tree produced by Bayesian inference, based on the amino acid sequences of the reverse transcriptase domain. The matrix was composed of the sequences of *X. tropicalis* and *X. laevis* DIRS-elements obtained from the Rebpase database, the copies retrieved from both genomes and diagnostic sequences from each DIRS superfamily. The posterior probabilities are indicated at the branches. Sequences of different superfamilies are highlighted in different colors: blue for *Ngaro-like*, green for *PAT-like*, and red for *DIRS-like*. The shades of each color also distinguish the sequences of *X. tropicalis* (dark) and *X. laevis* (light)

We recognized only two of the 95 *Xenopus* Rebpase DIRS families as belonging to the *Ngaro-like* superfamily, i.e., DIRS-53_XT and DIRS-54_XT. Sequences from the families DIRS-6A_XT, DIRS-13C_XT, DIRS-27A_XT, DIRS-35_XT, DIRS-42_XT, DIRS-3_XL, DIRS-10_XL, DIRS-13_XL, DIRS-15_XL, DIRS-18_XL, and DIRS-19_XL were not included in the tree because the RT domain was too short, but all these elements present a *DIRS-like* terminal repeat pattern.

The sequences of both superfamilies were analyzed, and a high level of congruence was found in the sequence structure in comparison with the DIRS families described in vertebrates (Goodwin and Poulter, 2004) (Figure 2). These findings will be discussed below.

**Fig. 2.**
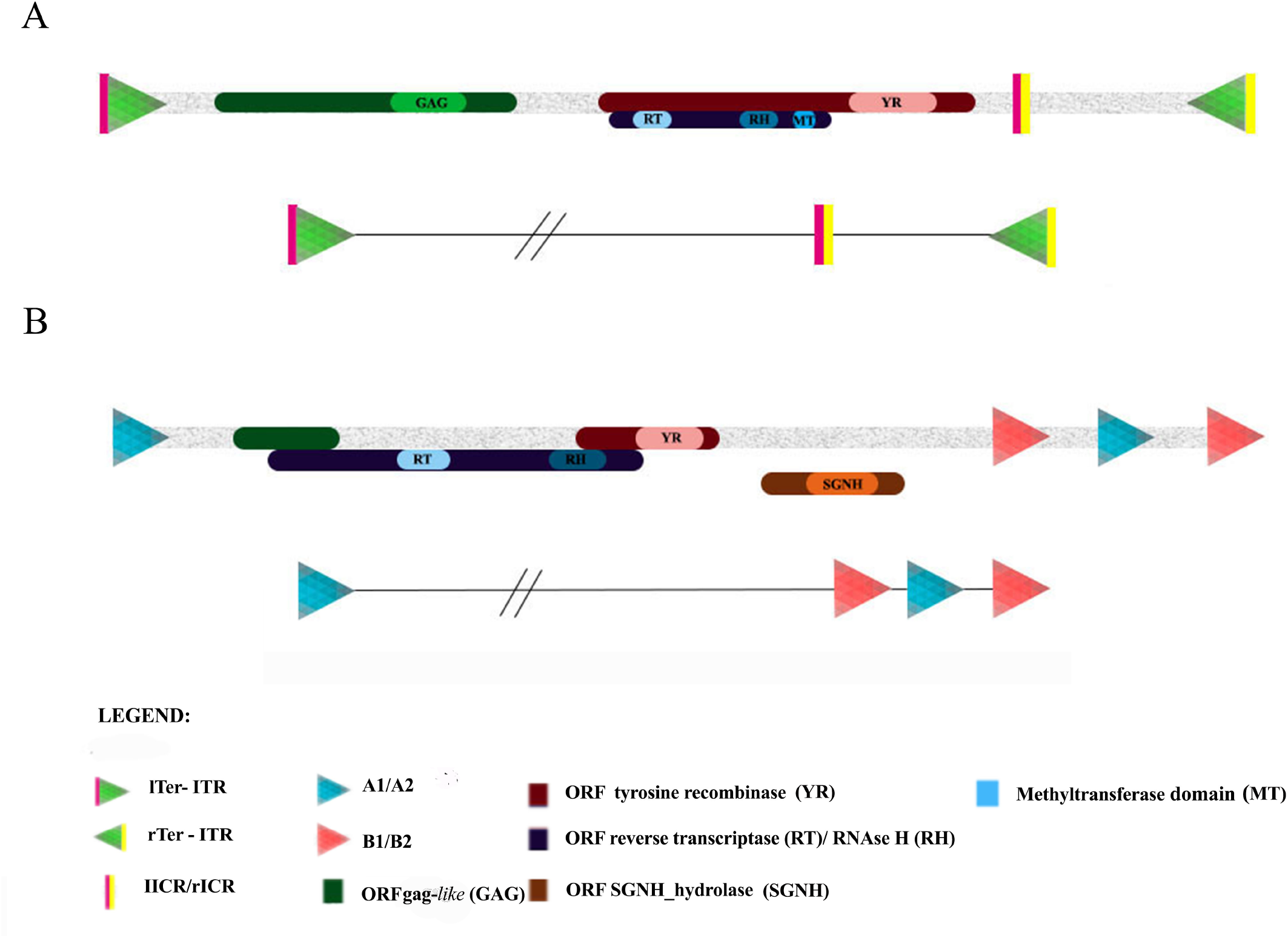
Schematic structure of the potentially complete *DIRS-like* and *Ngaro-like* retroelements of the *Xenopus tropicalis* genome. (A) Representation of the *X. tropicalis DIRS-like* elements, based on the Rebpase consensus sequences of the DIRS-37_XT, that contain three ORFs, conserved domains (gag, RT/RH/MT, and YR) and non-coding portions: inverted terminal repeats (ITRs) and internal complementary regions (ICRs). The expanded scheme of the terminal regions is shown in the lower plot. (B) Representation of the *X. tropicalis Ngaro-like* elements based on the Rebpase consensus sequences of the DIRS-53_XT that contain four ORFs, conserved domains (RT/RH, YR, and SGNH) and split direct repeats (SDRs)

### DIRS-like families

The consensus sequences of the *DIRS-like* superfamily found in the *X. tropicalis* and *X. laevis* genomes range from 4146 base pairs (bp) in DIRS-33_XT to 6224 bp in DIRS-2A_XT. We recognized three ORFs in almost all the families with several levels of overlap, involving primarily ORF2 and ORF3 (see Table 1 and Supplementary Table 1, for more details).

The ORF1 encodes a gag-like protein and a LAP2alpha domain (approximately 650 amino acids) was predicted in all the families. The ORF2 corresponds to the RT and RH domains, with around 120 and 356 aa, respectively. A deoxy-adenosine methylase (DAM/MT) domain of around 284 aa was also observed in the ORF2 of some of the families evaluated here (DIRS-9A_XT, DIRS-9_XT, DIRS-10_XT, DIRS-16_XT, DIRS-22A_XT, DIRS-22_XT, DIRS-25_XT, DIRS-26A_XT, DIRS-26_XT, DIRS-27A_XT, DIRS-27_XT, DIRS-29_XT, DIRS-31_XT. DIRS-32_XT, DIRS-34A_XT, DIRS-34_XT, DIRS-35_XT, DIRS-37A_XT, DIRS-37_XT, DIRS-38_XT, DIRS-41_XT, DIRS-41A_XT, DIRS-42_XT, DIRS-46_XT, DIRS-50_XT, DIRS-51_XT,DIRS-52_XT, DIRS-1_XL, DIRS-3_XL, DIRS-4_XL, DIRS-5_XL, DIRS-6_XL, DIRS-7_XL, DIRS-8_XL, DIRS-9_XL, DIRS-11_XL, DIRS-13_XL and DIRS-16_XL). The ORF3 encodes the YR protein with a conserved DNA_BRE_C domain of around 584 aa (Supplementary Table 1).

The *DIRS-like* elements have 5’ and 3’ ITRs and an ICR region (Figure 2A), and this pattern of repetition was found in almost all the *Xenopus DIRS-like* families evaluated here. The ITRs have approximately 120 bps and present a few nucleotide substitutions or *indels* between the left ITR (lITR) and the right ITR (rITR) (Figure 2A). The ICR is composed of two short sequences (lICR and rICR) which are complementary to the 5’ (lTer) and the 3’ (rTer) ends of the element (Figure 2A). The ICR and ITR sequences overlap slightly in most families.

Overall, 36 of the 75 Rebpase *DIRS-like* families of *X. tropicalis* present some level of degeneration in the molecular structure of the terminal repeats and/or ORFs domains (Table 1). In *X. laevis*, 13 of the 20 families present premature interruptions in the ORFs or incomplete repeats (Table 1), which indicates a high level of degeneration in these families (Supplementary Table 1). Concerning the EST data, we observed that most families present transcripts (55 families from *X. tropicalis* and 12 families from *X. laevis*) (Table 1). The *DIRS-like* families of both genomes have characteristic thymine trinucleotides (i.e., “TTT”) in both their 5’ and 3’ ends.

The RT sequence tree highlights the high level of diversity of the families of the *DIRS-like* clade in *Xenopus* (Figure 1). The diagnostic *DIRS-like* sequences (DIRS-1_ACar and DIRS-5_CBP) were recovered in an internal clade of the *Xenopus* sequences, which indicates that a number of *DIRS-like* families were already present in the ancestor of these species and thus contributed to the high current diversity of families.

As the Rebpase nomenclature of the families established for a species follows the order of their description (Kapitonov and Jurka, 2008; Bao et al. 2009), the evolutionary relationships among the families must be interpreted based on their phylogenetic relationships in the sequence trees, rather than their nomenclature in the databases. For example, DIRS-4_XT is not closely related to DIRS-4_XL, whereas DIRS-2_XL and DIRS-50_XT have a very close relationship. We recovered families of *DIRS-like* that were shared between the two species, such as DIRS-29_XT with DIRS-17_XL + DIRS-14_XL, DIRS-2_XL and DIRS-50_XT, DIRS-6_XL + DIRS-11_XL with DIRS-52_XT + DIRS-16_XT, and DIRS-41_XT + DIRS-41A_XT with DIRS-9_XL + DIRS-16_XL.

We also observed marked species-specific structuring in the sequence tree, recovering subclades that grouped families only from *X. tropicalis* or *X. laevis*. This indicates that many of the families may have originated after the separation of the two species, in particular in *X. tropicalis*. The landscape profile observed in each genome further reinforces this conclusion (Figure 3). The *X. laevis* landscape presents two broad ancient waves of invasion (of 60-90 million years ago - mya and 110-135 mya), whereas, in the *X. tropicalis* genome, there is a peak of very recent amplification, which occurred less than 3.2 mya. The most recent families of the *X. laevis* genome are found in smaller proportions, approximately 0.003% of the genome. Although the diversity of the *DIRS-like* families is much lower in *X. laevis*, they make up a larger proportion of the genome (0.8%) than in *X. tropicalis* (0.5%).

**Fig. 3.**
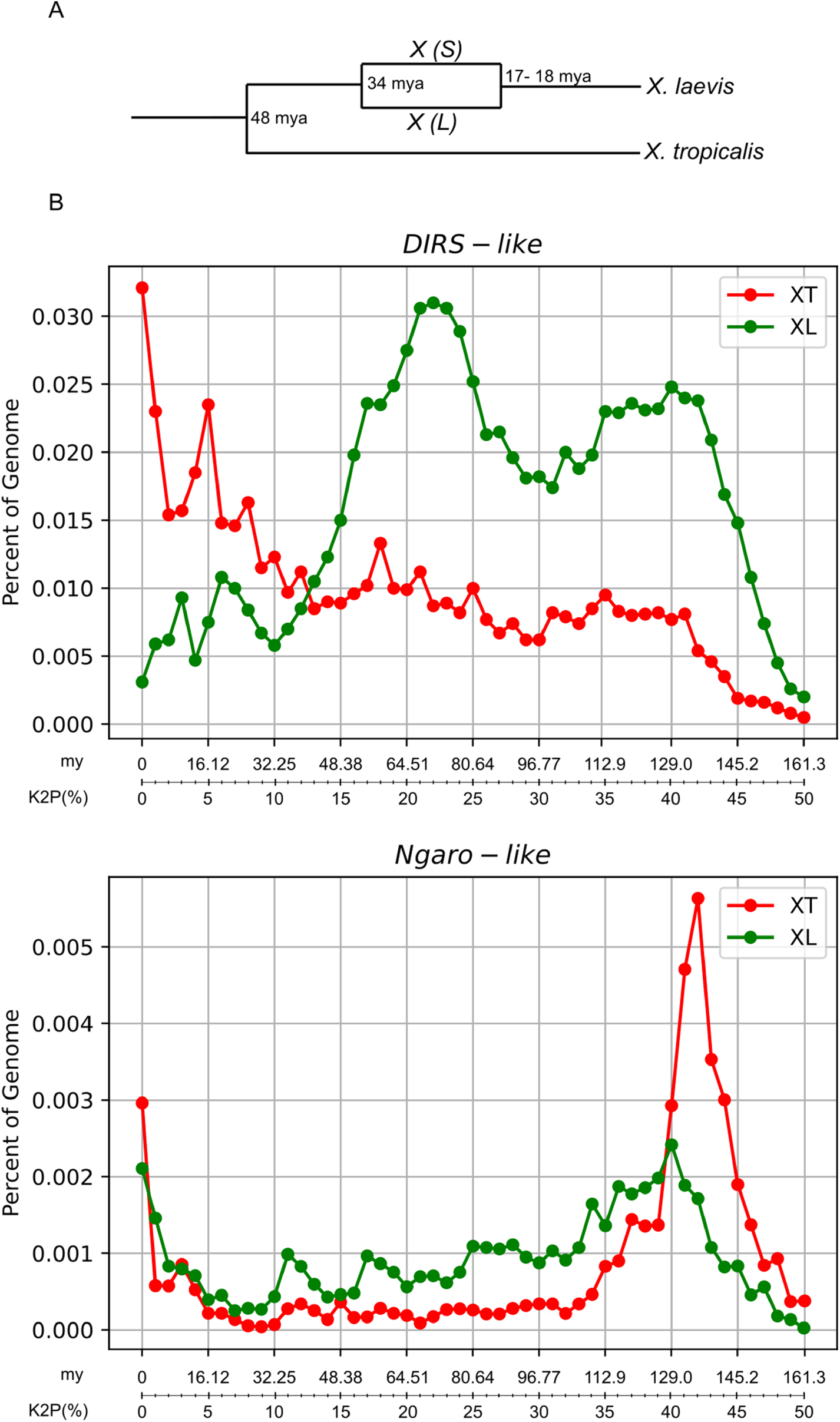
Evolution of *Xenopus* species and evolutionary dynamics of DIRS elements. A) Evolutionary events as estimated by Session et al (2016). The speciation of *X. tropicalis* and the ancestor of *X. laevis* at 48 mya, the speciation of the L and S progenitors at 34 mya, and the hybridization around 17-18 mya. B) The graphs show the divergence of *DIRS-like* (above) and *Ngaro-like* (below) copies mapped in the genomes of *X. tropicalis* (red) and *X. laevis* (green) with their consensus sequence expressed in Kimura-2-parameters distance and the corresponding time of divergence in million years (x-axis) plotted in relation to the proportion in the genome (y-axis).

From potentially active families, we chose to analyze the DIRS-37_XT copies in the genomes. In *X. tropicalis*, we can observe that most copies recovered (except copies 4 and 6) are putative functional presenting all ORFs and conserved ITRs and ICR. Copies 1 to 7 are the most closely related and grouped in the clade containing DIRS-37_XT, DIRS-37A_XT and DIRS-28_XT, but only the copies 1, 2 and 4 grouped with the query. The search has also recovered copies that grouped with more distant families, DIRS-31_XT (copies 8 and 9) and DIRS-3_XT (copy 10).

In *X. laevis*, the closest described families to DIRS-37_XT are DIRS-14_XL, DIRS-17_XL and DIRS-2_XL. All the copies recovered from the *X. laevis* genome present some level of degeneration, and due to this, copies 5 and 7 were not included in our phylogenetic inference. The nearest copy to the query is D37c1_XL, which was placed within a clade containing some *X. tropicalis* and no *X. laevis* families. This suggests that this sequence may be a degenerate copy of a *X. laevis* family that has yet to be established and was not successful in amplifying in this genome. Copies 2, 6, 8, and 9 grouped with DIRS-2_XL and DIRS-50_XT, while copies 3, 4, and 10 grouped with DIRS-14_XL, which is closely related to DIRS-17_XL and DIRS-29_XT. We found copies of the conserved elements in both subgenomes.

### Ngaro-like families

The *Ngaro-like* families DIRS-53_XT and DIRS-54_XT also present the three expected ORFs (encoding *gag*-like elements, RT/RH, and YR) with an additional ORF encoding a protein containing the SGNH_hydrolase domain. The reading frames of all the ORFs overlap (Figure 2B; Supplementary Table 1). The *Ngaro-like* elements are known to have a different type of terminal repeat, the SDRs (Goodwin and Poulter, 2004). This pattern can be observed in the DIRS-53_XT copy (Figure 2B). This element has type A1 and A2 SDRs of 236 bps and B1 and B2 types of 152 bps, with both repetitions being 100% identical. No SDRs were found in the DIRS-54_XT. Although none of the *X. laevis* Rebpase DIRS families grouped with *Ngaro-like* clade, our searches in the genome revealed the presence of sequences homologous to DIRS-53_XT and DIRS-54_XT, *Ngaro-like* transcripts were found for both species.

The copies of DIRS-53_XT and DIRS-54_XT recovered from *X. tropicalis* genome varied considerably in the degree of conservation of their sequences and structure (Supplementary Table 1), ranging from well-conserved copies to highly degenerate ones, due primarily to the loss of all or part of their 5’ and/or 3’ SDRs. Copies 1, 2, 4, 8, and 9 of DIRS-53_XT and copies 5 and 10 of DIRS-54_XT are potentially active.

In *X. laevis*, by contrast, we observed a high level of degeneration of the copies homologous to DIRS-53_XT/DIRS-54_XT, including a complete or partial loss of the 5’ and/or 3’ SDRs (Supplementary Table 1). These copies also present a high level of degeneration in the ORFs, including copies that lack the YR domain, hydrolase or RT domains (Supplementary Table 1). Of all copies recovered, only D54c5XL is potentially active.

The copies recovered from both species were included in the sequence tree (Figure 1), except those with a short RT domain (D53c2XT, D53c5XT, D53c9XT, D53c1XL, D53c2XL, D53c3XL, D53c4XL, D53c6XL, D53c7XL, D53c10XL, D54c1XL, D54c3XL, D54c6XL, D54c7XL, D54c9XL and D54c10XL). Clearly, DIRS-53_XT and DIRS-54_XT are very divergent (approximately 65% over only 600 bp of good alignment) and form two major groups of *Ngaro-like* sequences and some recovered copies did not group with the query. The DIRS-53 clade grouped copies retrieved with this query from *X. tropicalis* (copies 1, 3, 4, 6, 7, and 8) and copy 9 from *X. laevis*. The D54c5XL and D54c8XT copies also grouped with these sequences. This result suggests that this family is undergoing a process of degeneration in *X. laevis*, while it is still active in *X. tropicalis*.

The DIRS-54 clade contains the canonical element from the Rebpase, together with seven copies (1, 2, 3, 4, 5, 6 and 10) from the *X. tropicalis* genome (Figure 1). While the SDRs are not present in the consensus canonical copy, they can be observed in copies 5 and 10, which indicates that this family is also potentially active in *X. tropicalis*. None of *X. laevis* copies are grouped in this clade. The remaining copies from both species grouped in separate small clades and the copy D53c5XL is branching out from the base of the *Ngaro-like* clade, all of which are putative degenerate copies, given that none of them have SDRs. These sequences probably represent degenerate copies of the DIRS-53 or DIRS-54 families.

Contrary to what we observed for *DIRS-like, Ngaro-like* elements have similar densities in *X. tropicalis* and *X. laevis* of genomes, representing 0.049% and 0.047%, respectively. The *Ngaro-like* landscape profile shows a recent amplification signal (less than 3.2 mya) in the *X. tropicalis* and *X. laevis* genomes. Additionally, in both species it is possible to observe a very ancient amplification wave, dating from 120-145 mya, that is more prominent in *X. tropicalis* graph.

## DISCUSSION

In this work, we presented a detailed analysis of the YR-containing elements (order DIRS) available in the Repbase23.11 for *X. tropicalis* and *X. laevis* classifying them as either into the *DIRS-like* or *Ngaro-like* superfamilies. We have provided a detailed description of the structural characteristics of these elements and found the specific molecular signature of each superfamily. *DIRS-like* and *Ngaro-like* elements present distinct diagnostic features in their 5’ and 3’ non-coding terminal regions, with the same structural elements already recorded in other metazoans, fungi, and protists (Goodwin and Poulter 2001; Goodwin and Poulter 2004; Poulter and Goodwin 2005; Goodwin and Poulter 2015).

Our detailed analysis of the elements corroborates what has been described for the superfamilies also concerning the ORFs and domains (Goodwin and Poulter, 2001; Poulter and Goodwin, 2005). Thus, based on their molecular structure and phylogenetic criteria, only DIRS-53_XT_and DIRS-54_XT belong to the *Ngaro-like* superfamily, and all others are *DIRS-like*. We also identified conserved thymine trinucleotides (“TTT”) at the 5’ and 3’ ends in the complete copies of the *DIRS-like* elements of the genomes. This may be a general pattern of the *DIRS-like* elements found in terminal regions, which may be essential for transposing these elements (Malicki et al 2020). A similar terminal signature has also been described in the case of the DrDIRS1 element of *Tribolium castaneum*, which has either the trinucleotides ‘‘GTT’’ or dinucleotides “AA” (Goodwin et al. 2004). Given this, the recognition of the similarities in the molecular structure of these elements from the two species would also contribute to the assessment of the diversity and evolutionary history of these YR retrotransposons in other anuran genomes.

We found a much greater diversity and proportion of elements of the *DIRS-like* superfamily, in comparison with the *Ngaro-like* elements, in both genomes. *DIRS-like* and *Ngaro-like* elements are widely distributed in a number of metazoan groups, including fish, amphibians, reptiles, and in some fungus (Ruiz-Peres et al, 1996) and protists (Goodwin and Poulter, 2001; Poulter and Goodwin, 2005). Although there is no clear report of DIRS elements in some groups of species, we believe that the distribution of these retrotransposons is underestimated since they are frequently classified in the amount of the LTR retrotransposons group in the annotation of repetitive sequences based on the RepeatMasker tool.

The richness of the *DIRS-like* elements described in the present study had already been reported for the *X. tropicalis* genome (Hellsten et al. 2010; Piednoël et al. 2011) and reflects the importance of this repetitive element for its evolution. We recorded a much higher number of families of *DIRS-like* elements in *X. tropicalis* (81) than in the fish *Danio rerio* (12 families) or the reptile *Anolis carolinensis*, which has 42 families (Piednoël et al. 2011). We also found a large number of families are structurally complete in *X. tropicalis* and *X. laevis*. Thus, it is relevant to compare the landscape pattern of these superfamilies in both genomes since these species have undergone distinct evolutionary processes after the split. Session et al (2016) date the speciation of *X. tropicalis* and the *X. laevis* ancestor at around 48 mya. The *X. laevis* had an allotetraploid origin (around 17-18 mya) from two extant diploid progenitors separated at around 34 mya, and currently has two homoeologous subgenomes (L and S). The L and S subgenomes have undergone profound intragenomic diversification, which is compatible with the absence of recombination between the homeologous chromosome pairs of each subgenome since the allotetraploidization event (Session et al., 2016). The S subgenome has undergone intrachromosomal rearrangements and extensive small-scale deletions that resulted in the reduction of the length of the S chromosomes in comparison with their L homeologs (Sessions et al 2016).

If we consider the divergence time and evolutionary rates estimated by Session et al. (2016) as the most realistic scenario of the evolutionary history of *Xenopus* genus, our time-scale estimate of DIRS evolution in *X. tropicalis* and *X. laevis* genomes showed different patterns in both genomes. For *DIRS-like* it is possible to suggest that two ancient waves of amplification have occurred in the ancestor of both species. These ancient waves were less clear in *X. tropicalis* genome suggesting a higher turnover of degenerate copies in *X. tropicalis* than in *X. laevis* after the separation of these species.

Independent waves of *DIRS-like* amplification have also occurred after the separation of both species. It is possible to observe two more recent and modest amplification waves of *DIRS-like* in *X. laevis*, one estimated around 19 mya, probably in one or both ancestor genomes, and a more recent wave has occurred after the allotetraploidization, around 9.6 mya. This is consistent with the observation that some families are still potentially active in *X. laevis*. In *X. tropicalis* the amplification waves were more prominent occurring around 16 mya and less than 3.2 mya. The existence of potentially active copies of *DIRS-like* identified here is in agreement with the evidence of transcriptional activity detected in the transcriptome data on *X. tropicalis* (Poulter and Butler, 2015) and the searches performed by us in the EST libraries of both species. The recent burst events in both species could also have contributed to the diversification of families in each genome, primarily in *X. tropicalis*, which is consistent with the strong species-specific clustering seen in the sequence tree.

These two types of evidence (landscape pattern and phylogenetic) highlight the relative success of the *DIRS-like* elements in the *Xenopus* genomes in comparison with the *Ngaro-like* superfamily that presents very low diversity and quantity. *Ngaro-like* had a very ancient amplification wave followed by a long period of senescence. Despite *Ngaro-like* elements have failed to increase copy number and diversify in *Xenopus*, some potentially active copies are found in both genomes what is consistent with the very recent amplification wave (less than 3.2 Mya) seen in both species. The recent amplification indicates that *Ngaro-like* copies were maintained as active and somehow silenced for a long period of *Xenopus* evolution or were recently reactivated or reintroduced in these genomes. It is not clear, however, the reason why the *Ngaro-like* did not achieve the same success as *DIRS-like*, but it could be related to the known differences in the transposition mechanism (Poulter and Goodwin, 2005) of both superfamilies or to a possible variation in silencing efficiency by the host, while it could also be explained by chance. Still, evidence of the accumulation of DIRS-1 element on centromeres of the *D. discoideum* genome (Dubin et al. 2010; Malicki et al., 2017) could indicate their role during the centromeric heterochromatin biogenesis in this genome and open new perspectives to future evaluation about their biological significance in their success in *Xenopus* genomes.

The presence of very ancient and senescent DIRS copies in both genomes is consistent with “TE cycle life” of the genome (Kidwell and Lisch, 2001), in which ancient mobile elements may lose their autonomy and no amplifications occur, with the nucleotide sequences losing their identity, with the senescent elements eventually being deleted or becoming completely divergent. When we discriminated the *DIRS-like* and *Ngaro-like* families exclusive to each *X. laevis* subgenome, we observed a much greater diversity of families and copies in the L subgenome in comparison with the S subgenome (Supplementary Table 1). The identification of families exclusive of each subgenome in *X. laevis* was reported to Miniature Inverted-repeat Transposable Elements (MITE) Xl-TpL_Harb and Xl-TpS_Harb, exclusive from subgenome L and S, respectively and also support the absence of recombination between the homeologous chromosome pairs of each subgenome since the allotetrapoidization event (Session et al. 2016). The staggered pattern of superfamily diversity observed here (*X. tropicalis* > *X. laevis* L subgenome > *X. laevis* S subgenome) appears to reflect the same evolutionary pattern observed in other protein-coding genes, repetitive DNAs, and the rates of tandem duplications observed in each genome or subgenome (Session et al. 2016). However, as we found sequences in *X. laevis* from families non-reported in the Rebpase, we cannot discard the possibility that the diversity of DIRS in this species is somewhat larger.

Our findings also point to differential evolutionary dynamics of the YR retrotransposons of the *DIRS-like* order in the diploid (*X. tropicalis*) and allopolyploid *(X. laevis)* species studied here. The intragenomic behavior of the transposable elements depends on the balance between repression and expression, due to the need to avoid a large number of copies becoming a disadvantage for the genome (Bourque et al., 2018). The conservation of the molecular structure of these elements is related directly to these genetic mechanisms, which determine either an increase or loss of TE diversity, depending on the repertoire of TEs, during the genomic evolution of each lineage.

## Acknowledgements

We thank the Coordenação de Aperfeiçoamento de Pessoal de Nível Superior (CAPES/PROAP – Finance Code 001) for the scholarships provided to CBG. We acknowledge Xenbase (www.xenbase.org) for staging, gene expression resources, and genomic reference material consulted throughout this work. We especially thank Luciana Bolsoni Lourenço, Maríndia Deprá and Michelle Orane Schemberger for valuable comments on a previous version of the manuscript.

